# R/qtlcharts: interactive graphics for quantitative trait locus mapping

**DOI:** 10.1101/011437

**Authors:** Karl W. Broman

**Affiliations:** Departments of Biostatistics and Medical Informatics, University of Wisconsin–Madison, Madison, Wisconsin 53706

**Keywords:** QTL, data visualization, software, D3

## Abstract

Every data visualization can be improved with some level of interactivity. Interactive graphics hold particular promise for the exploration of high-dimensional data. R/qtlcharts is an R package to create interactive graphics for experiments to map quantitative trait loci (QTL; genetic loci that influence quantitative traits). R/qtlcharts serves as a companion to the R/qtl package, providing interactive versions of R/qtl’s static graphs, as well as additional interactive graphs for the exploration of high-dimensional genotype and phenotype data.

---

Interactive graphics have enormous value for the exploration of high-dimensional genetic data. Visualizations of high-dimensional data must be a compressed summary, but with interactive visualizations, features may be linked to underlying details or to different views of the data. Moreover, interactive graphs offer the ability to zoom into dense figures, and sets of linked graphic panels offer greater opportunity to make connections across diverse data types.

Interactive data visualization has a long history. For example, an early innovation, brushed scatterplots, is due to Becker and Cleveland (1987). While numerous tools for interactive data visualization have been developed, for example Mondrian (Theus and Urbanek 2008, http://www.theusrus.de/Mondrian) and SpotFire (http://spotfire.tibco.com/), until recently interactive visualization has been a specialized craft, and the tools have not been widely adopted as part of routine data analysis. But there has been a recent expansion in the general use and development of complex and rich interactive graphics tools, motivated in part by the JavaScript library D3 (Bostock *et al.* 2011, http://d3js.org), the development of HTML5 and scalable vector graphics (SVG), and the power of modern web browsers. Moreover, as these graphical tools are deployed as web pages, they are immediately accessible to users.

R/qtlcharts is add-on package for the general statistical software R (R Core Team 2013) that provides interactive data visualizations for quantitative trait locus (QTL) mapping experiments. R/qtlcharts is a companion to R/qtl (Broman *et al.* 2003), providing interactive versions of R/qtl’s basic static graphics and additional interactive visualizations for high-dimensional genotype and phenotype data.

For example, Figure 1 contains a static view of an interactive visualization of the results of QTL analysis with a phenotype measured over time. The data are from Moore *et al.* (2013).

**Figure 1:**
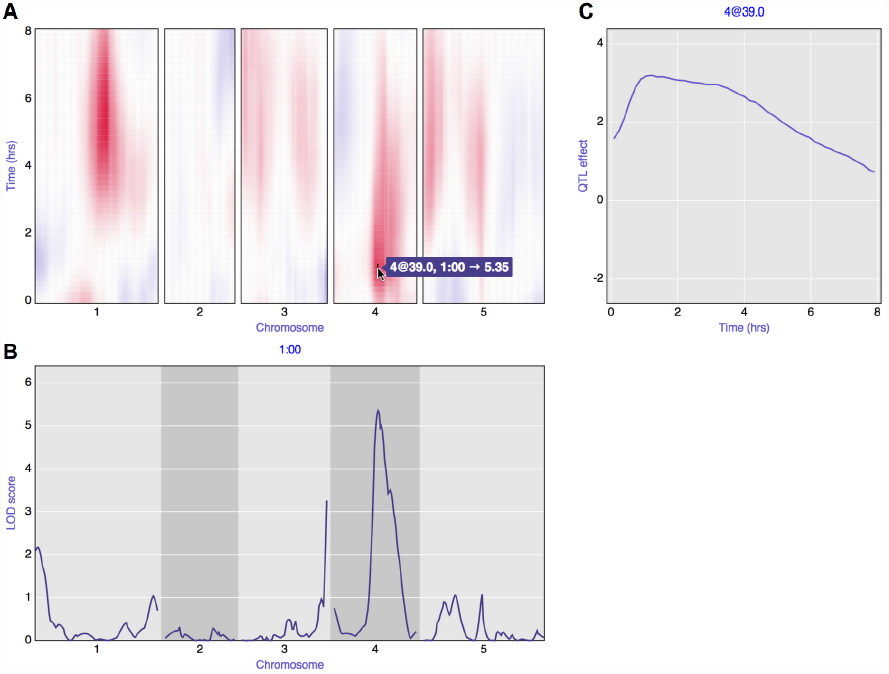
A static view of an interactive graph for QTL analysis with a phenotype measured over time, with data from Moore *et al.* (2013) concerning root gravitropism in Arabidopsis recombinant inbred lines (RIL), Ler*×*Cvi. **A**: Heat map of signed LOD scores for single-QTL analysis at each individual time point.Red indicates that RIL with the Cvi allele have a larger average phenotype; blue indicates that RIL with the Ler allele have a larger average phenotype. Panel A is linked to panels B and C: when hovering over the heat map, the LOD curves for the corresponding time point are shown below (**B**), and the estimated QTL effect, as a function of time, is shown to the right (**C**). For the interactive version of this figure, see http://kbroman.org/qtlcharts/example.

Figure 1A contains a heat map of the LOD scores from a single-QTL genome scan, considering each time point individually. Hovering over the heat map drives the contents of the other two panels and also reveals the LOD score values through a tool-tip. Figure 1B displays the LOD curves across the genome for the selected time point. Figure 1C displays the estimated effect, across time, of the putative QTL at the selected position.

The interactive versions of R/qtl’s static graphs provide considerable value to the user, enabling a deeper and more rapid exploration of QTL mapping results. The LOD curves from a single-QTL genome scan are linked to a panel displaying the association between genotype and phenotype at putative QTL. This lowers the barrier to inspection of QTL effects. One should not rely solely on test statistics like the LOD score, but with the extra effort required to generate plots of QTL effects in R/qtl, such effect plots are sometimes skipped. The interactive graph of genome scan results can also serve as important tool for teaching: the easy connection between LOD scores and estimated QTL effects can help students to develop a better understanding of QTL analysis results.

A static heat map with the results of a two-dimensional, two-QTL genome scan can be difficult to interpret. In the interactive version, hovering over the heat map with a mouse reveals the corresponding positions and LOD scores, and mouse click generates plots of the genotype-phenotype relationship for the pair of putative QTL, as well as plots of cross-sectional slices through the two-dimensional image. This interactivity eases the identification of both interesting QTL effects as well as artifacts (such as apparent epistatic interactions that are driven by a few individuals with outlying phenotype values).

One final example: in a static graph of a genetic marker map, the marker names are often omitted, as they otherwise clutter the figure. In the interactive version produced by R/qtlcharts, the marker names and locations are revealed when hovering over the positions, and there is also a search box for identifying the position of a specified marker.

The core of R/qtlcharts is a set of modular, reusable graphic panels, based on the D3 JavaScript library. Examples of these panels are displayed in Figure 2 and include heat maps, LOD curve plots, and scatterplots. Multiple panels are combined into higher-level interactive charts with links among the panels (as in Figure 1).

**Figure 2:**
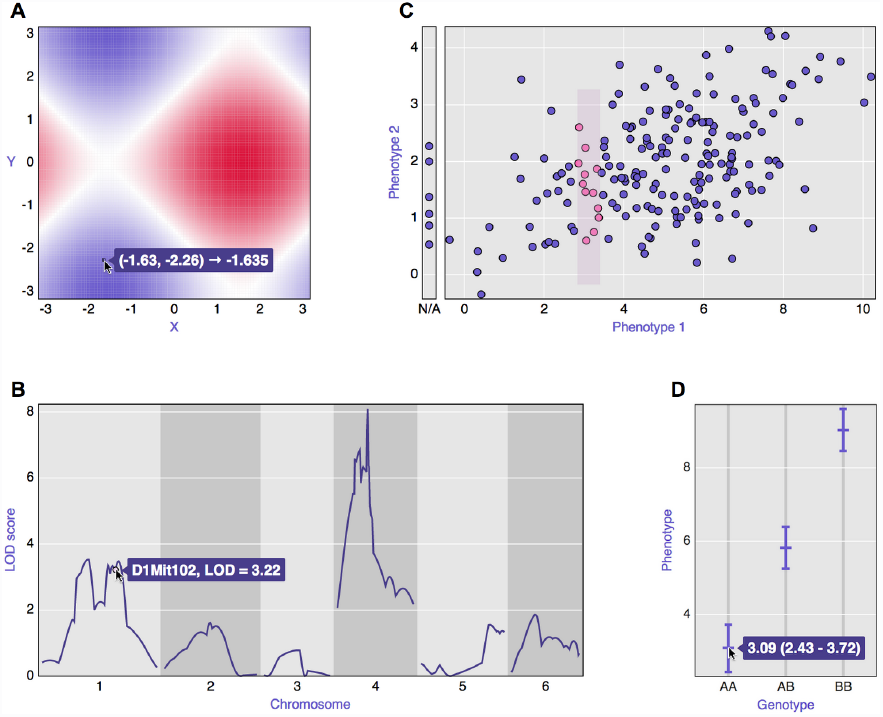
Examples of the basic panels that form the core of R/qtlcharts. **A**: heat map, **B**: LOD curves, **C**: scatterplot, and **D**: a set of confidence intervals.

With R/qtlcharts, a single line of R code can generate such an interactive graph: an HTML file is created and opened in a web browser. The file can be self-contained and so easily shared with collaborators.

The interactive charts can also be included within R Markdown documents (http://rmarkdown.rstudio.com). R Markdown is a variant of the light-weight markup language, Markdown (http://daringfireball.net/projects/markdown), with embedded chunks of R code. An R Markdown document is converted, via knitr (Xie 2013, http://yihui.name/knitr), to an HTML file with the R code chunks replaced by their results (including, for example, figures). This is an example of literate programming (Knuth 1984), a key tool for ensuring that research results are reproducible.

The interactive graphics in R/qtlcharts are written in CoffeeScript (http://coffeescript.org), which is translated to JavaScript but is a more pleasant language for programming. Tool tips are implemented with d3-tip (https://github.com/Caged/d3-tip).

Examples and documentation, including installation instructions, are available at the R/qtlcharts website, http://kbroman.org/qtlcharts. The code is released under the MIT license and is available at GitHub, http://github.com/kbroman/qtlcharts.

## Ackwnoledgements

Śaunak Sen generously provided comments for improvement of the manuscript. This work was supported in part by National Institutes of Health grant R01GM074244.

